# Immune Evasion, Infectivity, and Fusogenicity of SARS-CoV-2 Omicron BA.2.86 and FLip Variants

**DOI:** 10.1101/2023.09.11.557206

**Authors:** Panke Qu, Kai Xu, Julia N. Faraone, Negin Goodarzi, Yi-Min Zheng, Claire Carlin, Joseph S. Bednash, Jeffrey C. Horowitz, Rama K. Mallampalli, Linda J. Saif, Eugene M. Oltz, Daniel Jones, Richard J. Gumina, Shan-Lu Liu

## Abstract

Evolution of SARS-CoV-2 requires the reassessment of current vaccine measures. Here, we characterized BA.2.86 and the XBB-lineage variant FLip by investigating their neutralization alongside D614G, BA.1, BA.2, BA.4/5, XBB.1.5, and EG.5.1 by sera from 3-dose vaccinated and bivalent vaccinated healthcare workers, XBB.1.5-wave infected first responders, and monoclonal antibody (mAb) S309. We assessed the biology of the variant Spikes by measuring viral infectivity and membrane fusogenicity. BA.2.86 is less immune evasive compared to FLip and other XBB variants, consistent with antigenic distances. Importantly, distinct from XBB variants, mAb S309 was unable to neutralize BA.2.86, likely due to a D339H mutation based on modeling. BA.2.86 had relatively high fusogenicity and infectivity in CaLu-3 cells but low fusion and infectivity in 293T-ACE2 cells compared to some XBB variants, suggesting a potentially differences conformational stability of BA.2.86 Spike. Overall, our study underscores the importance of SARS-CoV-2 variant surveillance and the need for updated COVID-19 vaccines.

## Introduction

One of the biggest challenges faced throughout the COVID-19 pandemic is the speed with which the causative agent SARS-CoV-2 mutates^1^. The ongoing evolution of the virus has made it challenging to update and maintain current vaccination measures. This issue was exacerbated with the emergence of the Omicron BA.1 variant in late 2021, which is characterized by over 30 new mutations in Spike alone, as well as subsequent Omicron sublineages harboring additional mutations^1^. These mutations contributed to notable changes in the biology of the virus, including increased transmissibility^2^, decreased pathogenicity^2–4^, and marked immune evasion^5–11^. Immune evasion by these variants has reached a new threshold with the emergence of the recombinant XBB lineage of Omicron subvariants in early 2023, including XBB.1.5, XBB.1.16 and XBB.2.3. These variants exhibited dramatic escape of neutralizing antibodies stimulated through 3-dose vaccination that can be partially recovered through the administration of a bivalent mRNA booster^12–21^. The escape variants have has led to the decision by government regulators to include XBB Spikes in the newest versions of the mRNA vaccines this fall^22–24^.

Of current concern is a new variant, referred to as the “second generation BA.2”, named BA.2.86. BA.2.86, which was first detected in late July 2023 in Israel and Denmark^25,26^, and now has been documented in different parts of the world, including Australia, Canada, France, United Kingdom (U.K.), and the United States (U.S.). The Spike protein of BA.2.86 is characterized by more than 30 mutations relative to the predicted ancestral variant BA.2 and ∼35 mutations distinct from XBB.1.5^27^ (**Fig. 1A**). The number of mutations in Spike is reminiscent of the original Omicron BA.1 relative to previous variants of concern. Importantly, there have been several confirmed cases and detection of the variant in wastewater in some locations including the states of New York and Ohio in the U.S. The cases appear to be independent of each other, and many are individuals who have not traveled recently^28–34^, suggesting possible widespread dissemination of this variant. Of particular note is an outbreak in a U.K. care home that has so far resulted in at least 28 cases, demonstrating the variant’s ability to transmit in a close-contact setting^33^. These findings have led to the increased surveillance of BA.2.86 and its characterization as a “variant under monitoring” in the U.K and U.S^25,35^.

**Figure 1:**
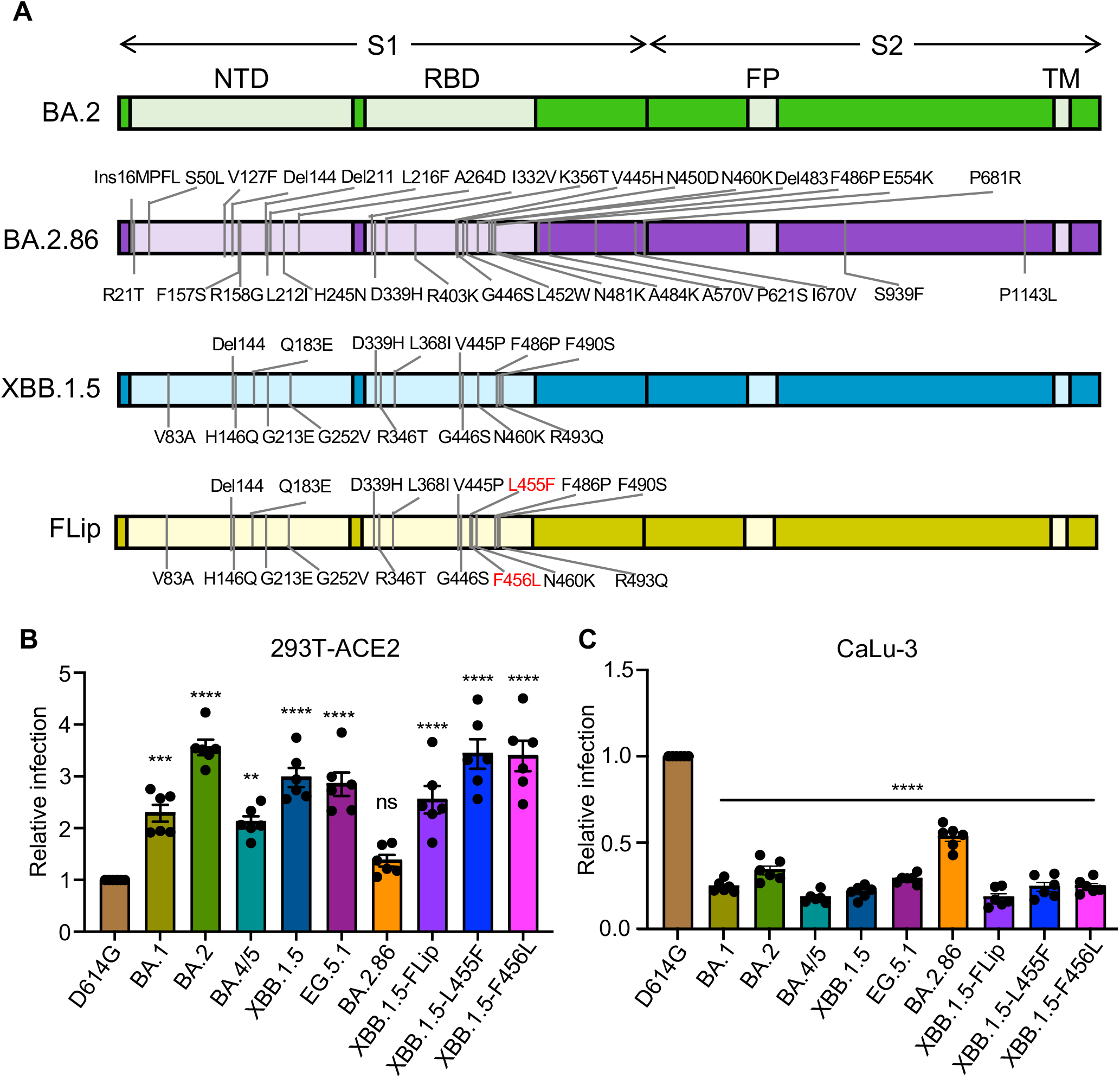
Infectivity of Omicron subvariants BA.2.86 and FLip. (**A**) Diagrams of the SARS-CoV-2 Omicron subvariants BA.2, BA.2.86, and XBB.1.5 Spikes. The location of specific mutations for BA.2.86 or XBB.1.5 relative to BA.2 in the N-Terminal Domain (NTD) or Receptor Binding Domain (RBD) of the S1 subunit, or in the domain between Fusion Peptide (FP) and Trans-membrane domain (TM) of the S2 subunit, or near the S1/S2 cleavage site is shown. (**B-C**) Infectivity of pseudotyped lentiviruses bearing each of the indicated Omicron subvariants Spike was determined in (**B**) HEK293T cells stably expressing human ACE2 (HEK293T-ACE2) or (**C**) human lung cell-derived epithelial CaLu-3 cells. Transfection efficiency and Spike protein expression were comparable among all groups, which is shown in Figure 5C. Bars in (**B-C**) represent means ± standard error from triplicates. Significance relative to D614G was analyzed by a one-way repeated measures ANOVA with Bonferroni’s multiple testing correction (n=6). P values are displayed as ns p > 0.05, **p < 0.01, ***p < 0.001, and ****p < 0.0001.

Given that BA.2.86 Spike is notably distinct from XBB.1.5, there is concern that current mRNA vaccines as well as the updated XBB.1.5 mRNA booster, will not effectively protect against BA.2.86^31,32^. Some initial studies have been performed to determine whether BA.2.86 may have growth advantages comparable to Omicron BA.1, particularly its ability to escape neutralizing antibodies. Deep mutational scanning analysis by the Bloom group revealed 17 mutations that have the potential to disrupt neutralizing antibody binding, largely concentrated around the N-terminal domain (NTD) and receptor binding domain (RBD) (**Fig. 1A**). Their data suggested that BA.2.86 will be about as immune evasive as XBB.1.5, but antigenically distinct from XBB lineage variants^32^. However, some recent data indicate that BA.2.86 is not as immune evasive as XBB.1.5 and other XBB variants. Hence, it is critical to understand whether current vaccination measures can still produce antibodies that effectively neutralize BA.2.86. Additionally, it is currently unknown whether BA.2.86 may exhibit growth advantages over other currently circulating Omicron variants, including EG.5.1 and the FLip variant, which contains the L455F and F456L mutation in the background of XBB.1.5^1^. In this study, we characterized neutralizing antibody titers against BA.2.86 alongside D614G, BA.1, BA.2, BA.4/5, XBB.1.5, EG.5.1, and FLip for bivalent vaccinated health care workers (HCWs) (n=14), monovalent 3-dose vaccinated HCWs (n=15), XBB.1.5-wave infected individuals (n=11), and monoclonal antibody S309 which has been shown to be effective against most Omicron variants including XBB 1.5 and EG.5.1^16,36–38^. We also characterize the biology of the BA.2.86 Spike by investigating pseudotyped viral infectivity, membrane fusogenicity, and Spike processing compared with other SARS-CoV-2 variants.

## Results

### Infectivity of BA.2.86 and Flip

First, we determined the infectivity of lentiviral pseudotypes bearing each of the SARS-CoV-2 Spikes of interest in HEK293T cells expressing human ACE2 (293T-ACE2) and in human lung adenocarcinoma cell line CaLu-3. In 293T-ACE2 cells, BA.2.86 did not exhibit a significant change in infectivity compared to D614G (1.4- fold increase; p > 0.05) but showed a 2.6-fold drop relative to BA.2 (p < 0.001) (**Fig. 1B**). Notably, the infectivity of BA.2.86 was 1.8∼2.1-fold lower than all Omicron variants including XBB.1.5 and EG.5.1. In contrast, the FLip variant exhibited a 2.5-fold and 1.8-fold increased titer compared to D614G (p < 0.0001) and BA.2.86 (p < 0.01), respectively, with a level comparable to XBB.1.5 and EG.5.1 (**Fig. 1B**). Both XBB.1.5-L455F and XBB.1.5-F456L contributed equally to the increased infectivity of FLip, with 3.4-fold increase, relative to D614G (p < 0.0001) (**Fig. 1B**).

In CaLu-3 cells, BA.2.86 exhibited significantly decreased infectivity relative to D614G (p < 0.0001), similar to all Omicron variants (**Fig. 1C**). Intriguingly, BA.2.86 showed a 1.9∼2.8-fold increase in infectivity compared to XBB.1.5, EG.5.1 and Flip (p < 0.0001). The FLip variant exhibited a 5.3-fold reduction in titer relative to D614G (p < 0.001), again more closely resembling other Omicron subvariants (p < 0.001), with both the XBB.1.5-L455F and XBB.1.5-F456L mutations (p < 0.0001) contributing to this phenotype (**Fig. 1C**). Overall, in comparison to earlier Omicron XBB subvariants, BA.2.86 appears to have a decreased infectivity in 293T-ACE2 cells yet increased infectivity in CaLu-3 cells. In contrast, the FLip variant follows the same trends of comparable infectivity to other XBB variants, including XBB.1.5 and EG.5.1, in both 293T-ACE2 and CaLu-3 cells.

### BA.2.86 is less resistant to neutralization by bivalent boosted sera compared to XBB.1.5, EG.5.1, and FLip

We determined the sensitivity of new Omicron variants BA.2.86 and FLip to neutralization by sera of a cohort of health care workers (HCWs) that received at least 2 doses of monovalent vaccine and 1 dose of bivalent mRNA booster (n = 14, **Table S1**). Consistent with what we have reported previously^16,36^, the neutralizing antibody (nAb) titers of these samples against Omicron subvariants were higher compared to the 3-dose monovalent vaccinated cohort (**Fig. 2A-D**)^16^. As might be expected, BA.2.86 exhibited reduced nAb titers, with 12.8-fold relative to D614G (p < 0.0001) and 11.7-fold compared to BA.2 (p < 0.0001), respectively (**Fig. 2A-B; Fig. S1A**). Strikingly, we observed a 1.7∼5.5-fold increased nAb titer against BA.2.86 compared to other recently emerged XBB variants, including XBB.1.5 (p >0.05), EG.5.1 (p < 0.01), and FLip (p < 0.0001). The latter 3 variants had 21.9-fold, 36.6-fold and 70.9-fold reductions, respectively, in titer relative to D614G (p < 0.0001 for all 3 variants). Notably, FLip exhibited more nAb escape than its parental variant XBB.1.5, with a 3.2-fold reduction in titer (p < 0.0001) due to both the XBB.1.5-L455F and XBB.1.5-F456L mutations (2-fold for each, p < 0.01). Overall, bivalent vaccinated HCW sera neutralized BA.2.86 more efficiently than other XBB variants, while FLip exhibited much more pronounced nAb escape than other XBB variants.

**Figure 2:**
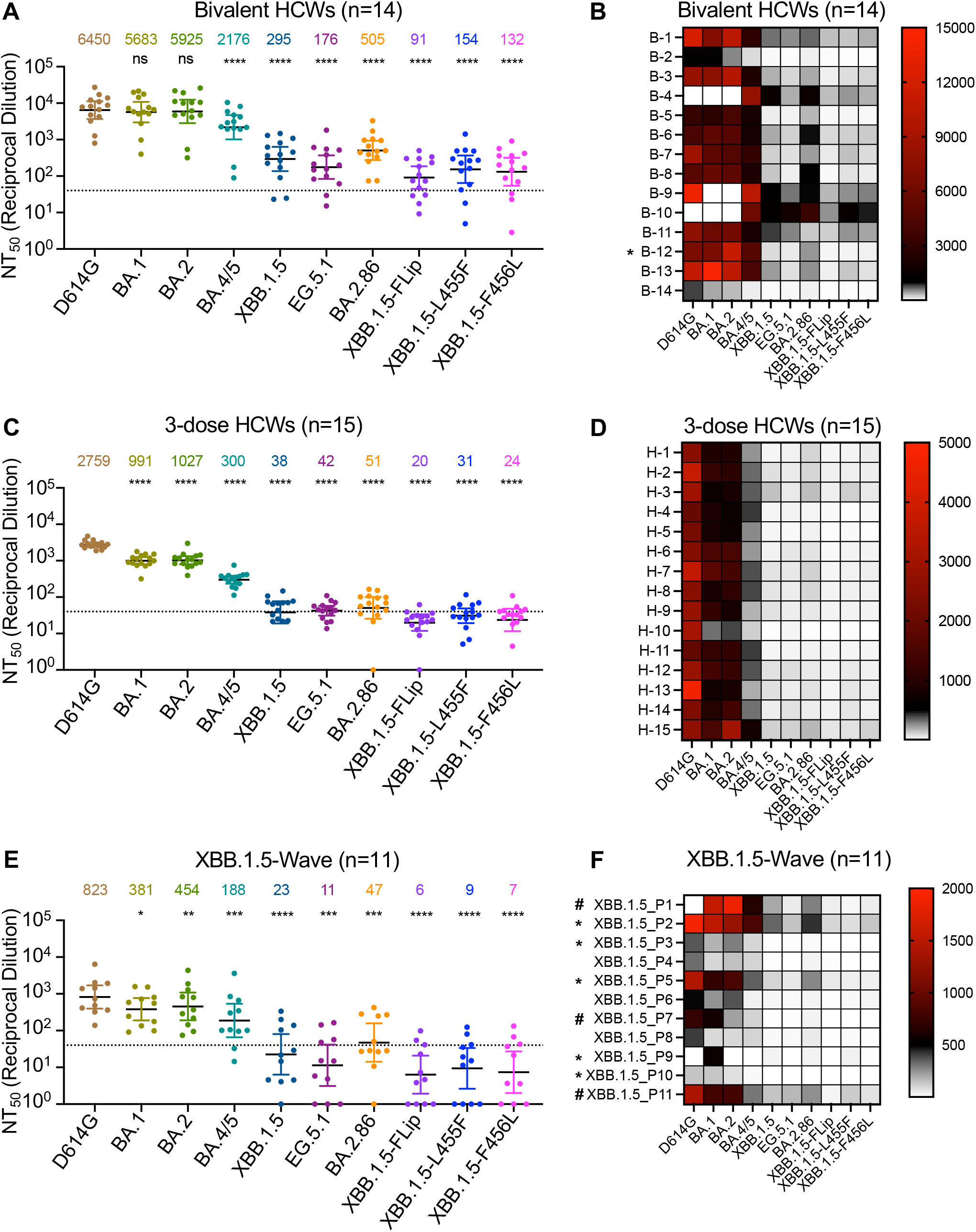
Neutralization of Omicron BA.2.86 and FLip subvariants by sera of monovalent or bivalent mRNA vaccinated health care workers (HCWs) and XBB.1.5-wave infection. Neutralizing antibody (NAb) titers were determined using lentiviruses containing the indicated Spike proteins with D614G as a control. All the NAb titers were compared against D614G. The three cohorts included sera from 14 HCWs who received 3 monovalent doses of mRNA vaccine and 1 dose of bivalent mRNA vaccine (n=14) (**A** and **B**), sera from 15 HCWs that received three doses of monovalent mRNA vaccine (n=15) (**C** and **D**), and sera from 11 SARS-CoV-2 infected first responders/household contacts or hospitalized patients who tested COVID-19 positive during the XBB1.5- wave of infection in Columbus, Ohio (**E** and **F**). Geometric mean NT_50_ values for each variant are shown on the top. Bars represent geometric means with 95% confidence intervals. Statistical significance was analyzed with log10 transformed NT_50_ values. Comparisons between multiple groups were performed using a one-way ANOVA with Bonferroni post-test. Dashed lines represent the threshold of detection, i.e., NT_50_=40. P values are shown as ns p > 0.05, *p < 0.05, **p < 0.01, ***p < 0.001, ****p < 0.0001. Heatmaps in (**B**, **D** and **F**) indicate NAb titers of each individual against each variant tested. Asterisk in (**B**) indicates that the person being COVID-19 positive within six months before the sera collection, asterisk in (**F**) indicates that the individuals who received 2 or 3-dose monovalent vaccines before infection, and number sign in (**F**) indicates that the individuals that received monovalent vaccines and bivalent vaccines.

### Neutralizing antibodies in 3-dose vaccinated sera are unable to neutralize BA.2.86 similar to XBB variants

We next examined the nAb titers in 3-dose mRNA vaccinated HCWs (n = 15) that have received at least 2 homologous doses of either Pfizer or Moderna monovalent mRNA vaccines (**Table S1**). Similar to XBB variants including XBB.1.5 and EG.5.1, BA.2.86 exhibited nAb titers around the limit of detection for the assay, i. e, NT_50_ = 40, with a 54.1-fold reduction compared to D614G (p < 0.0001) and a 20.1-fold reduction relative to its parental BA.2 (p < 0.0001), respectively (**Fig. 2C-D**). Notably, the FLip variant exhibited a more dramatic escape, with 138.0-fold and 51.4-fold reduced nAb titers relative to D614G and BA.2, respectively (p < 0.0001) (**Fig. 2C-D)**. This extent of nAb escape by FLip was largely comparable to its parental variant XBB.1.5, with NT_50_ values all falling below the limit of detection, and were due to both the XBB.1.5-L455F and XBB.1.5-F456L mutations (**Fig. 2C-D**). Overall, BA.2.86 and FLip variants exhibit marked escape of nAbs in 3-dose monovalent vaccinated sera, with titers near or below the limit of detection.

### XBB.1.5-wave breakthrough infections conferred almost no nAb against BA.2.86 and FLip variants

The final cohort we investigated were individuals who became infected during the XBB.1.5 wave in Columbus, Ohio (n = 11). Nasal swabs were performed to confirm COVID-19 positive status of 8 individuals and sequencing identified XBB.1.5 as the infecting variant; 3 samples were not sequence confirmed but collected after February, 2023 when XBB variants had become dominant in this area. Among these 11 samples, 8 were vaccinated, 3 of which received 3 doses of monovalent vaccine, 3 received at least 3 doses of monovalent and 1 dose bivalent booster, 2 received 2 doses of monovalent vaccine (**Table S1**). Overall, the nAb titers against all variants in this cohort were much lower than in the bivalent or monovalent-vaccinated cohorts, with NT_50_ below the limit of detection for all XBB variants (**Fig. 2E-F**). Of note, BA.2.86 exhibited an average of NT_50_ = 50, which was slightly above the limit of detection, i.e, NT_50_ = 40. The nAb titers against FLip were the lowest among all the variants examined (**Fig. 2E-F**). Importantly, 3 to 5 of the 11 individuals who had received at least 3-dose mRNA vaccine (**Table S1**) exhibited nAb titers above the limit of detection for FLip or BA.2.86 (**Fig. 2E-F, Fig. S1B**). In summary, while XBB.1.5-wave breakthrough infections confer limited if any neutralization against BA.2.86 and FLip, BA.2.86 still exhibits less nAb evasion compared to XBB variants in the XBB.1.5-convalescent cohort.

### Monoclonal antibody S309 neutralizes FLip but not BA.2.86

Monoclonal antibody (mAb) treatments have been crucial in the control during the early stages of the COVID-19 pandemic^39^, and remarkably one of the mAbs, i.e., S309, has been shown to neutralize all Omicron subvariants, including XBB.1.5, XBB.1.6, XBB.2.3 and EG.5.1^16,40,41^. Surprisingly, we found that S309 was unable to neutralize BA.2.86, with no inhibitory concentration at 50% (IC_50_) detectable. This was in stark contrast to FLip and other Omicron variants, which were efficiently neutralized by S309, with IC50 between 0.34 ± 0.13 (BA.1) µg/mL and 5.50 ± 0.75 (FLip) µg/mL (**Fig. 3A-B**). These results indicated that BA.2.86 is resistant to S309, which is distinct from other SARS-CoV-2 variants including XBB.1.5, EG.5.1 and FLip (see Discussion).

**Figure 3:**
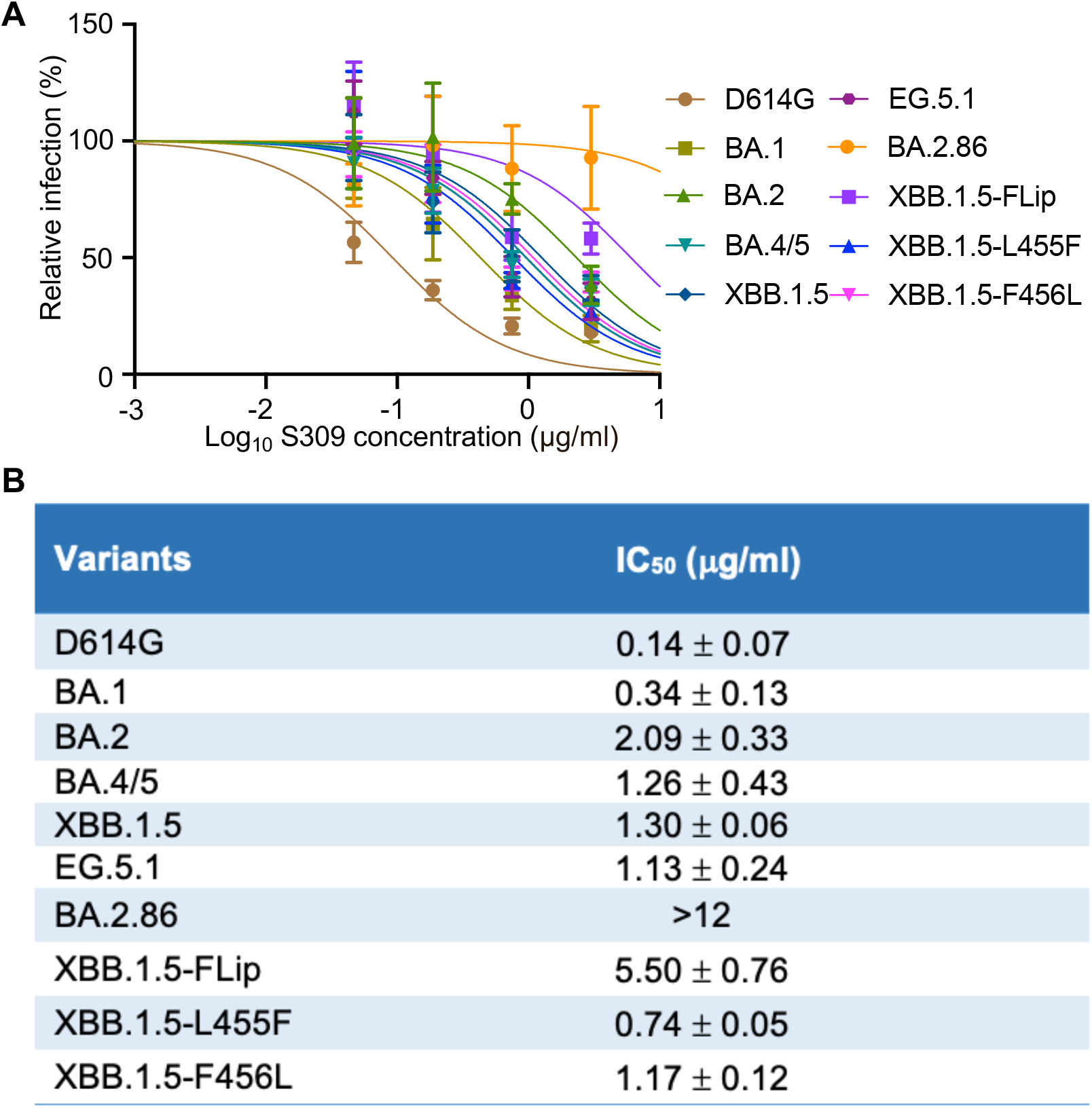
Neutralization of Omicron BA.2.86 and FLip subvariants by monoclonal antibody (mAb) S309. The lentiviral pseudotypes carrying each of the indicated variant Spike proteins were used to assess the effectiveness of mAb S309 in neutralizing BA.2.86, Flip and other variants. Representative plot curves are displayed (**A**) and the calculated IC_50_ values (means ± standard deviation) from two biological replicates are shown (**B**).

### BA.2.86 Spike has low fusogenicity in 293T-ACE2 cells, the activity of which is overcome in CaLu-3 cells

To understand the possible mechanisms underlying the differential infectivity of BA.2.86 and other Spikes in 293T-ACE2 and CaLu-3 cells, we investigated their ability to induce membrane fusion as well as Spike processing. For cell-cell fusion, we transfected effector 293T cells with Spike plasmid of interest plus GFP, and cocultured the effector cells with target cells, either 293T-ACE2 or CaLu-3 cells in parallel, with cell-cell fusion efficiency examined by imaging and quantified using the Leica X Applications Suite software. Similar to our previous results^5,11,20,42^, all Omicron variants exhibited markedly reduced cell-cell fusion compared to D614G (**Fig. 4A-B**). Notably, in contrast to XBB variants, especially XBB.1.5 and EG.5.1 which exhibited relatively high fusion activities, BA.2.86 showed a reduction in cell-cell fusion, with the level almost comparable to the ancestral BA.2/BA.1. This reduced fusion appeared consistent with the low infectivity/entry of BA.2.86 in 293T-ACE2 cells (**Fig. 1A**). Interestingly, we found that the low cell-cell fusion activity of BA.2.86 between 293T and 293T-ACE2 cells was rescued when 293T and CaLu-3 cells were cocultured, which showed increased fusion for BA.2.86 as compared to XBB.1.5. The level of fusion in CaLu-3 cells for BA.2.86 was almost comparable to that of FLip (**Fig. 4C-D**).

**Figure 4:**
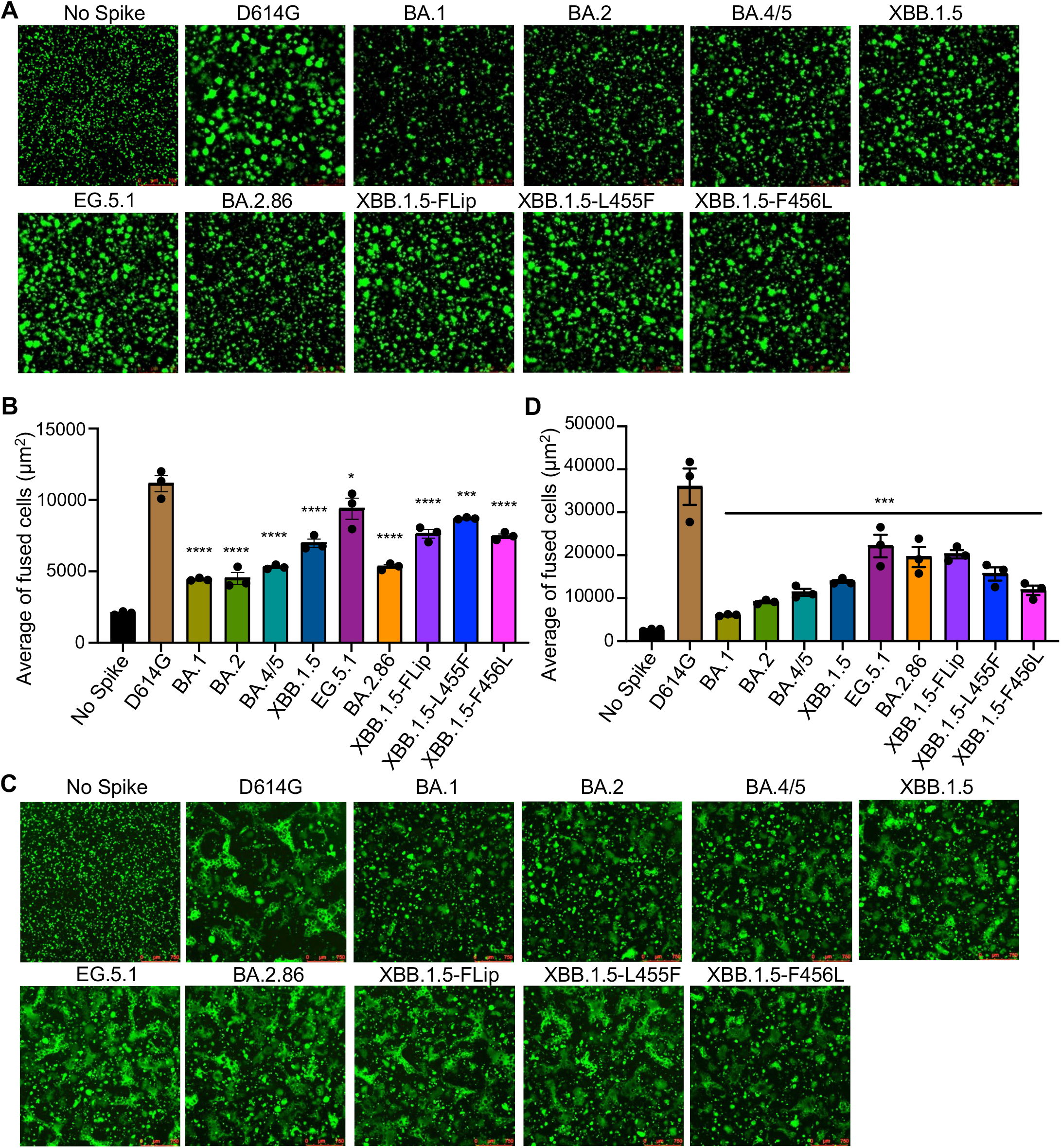
Cell-cell fusion of Omicron BA.2.86 and FLip subvariants. HEK293T cells were cotransfected with the indicated Spikes of interest and GFP plasmids, and were cocultured with HEK29T cells overexpressing human ACE2 (293T-ACE2) (**A-B**) or human lung epithelial CaLu-3 cells (**C-D**) for 24 hours. Cell-cell fusion was imaged and GFP areas of fused cells were quantified (see Methods). D614G and no S were included as positive and negative control, respectively. Comparisons in extents of cell-cell fusion for each Omicron subvariant were made against D614G. Bars in (**B** and **D**) represent means ± standard error. Dots represent three images from two biological replicates. Statistical significance relative to D614G was determined using a one-way repeated measures ANOVA with Bonferroni’s multiple testing correction (n=3). P values are displayed as ns p > 0.05, *p <0.05, ***p < 0.001, and ****p < 0.0001.

We examined the expression levels of Spike proteins on the plasma membrane of transiently transfected cells by performing flow cytometry using a polyclonal antibody against S1. We observed approximately similar levels of expression for Spikes, with BA.2.86, XBB.1.5 and D614G being approximately 50% lower than other variants (**Fig. 5A-B**). In addition, we determined the Spike processing of these variants in 293T cells producing the pseudotyped viruses. We found a decreased level of BA.2.86 Spike processing as compared to XBB variants including XBB.1.5, EG.5.1 and FLip, all of which showed a higher level of Spike processing relative to D614G, BA.1 and BA.2 based on the calculated ratios of S2/S and S1/S (**Fig. 5C**).

**Figure 5:**
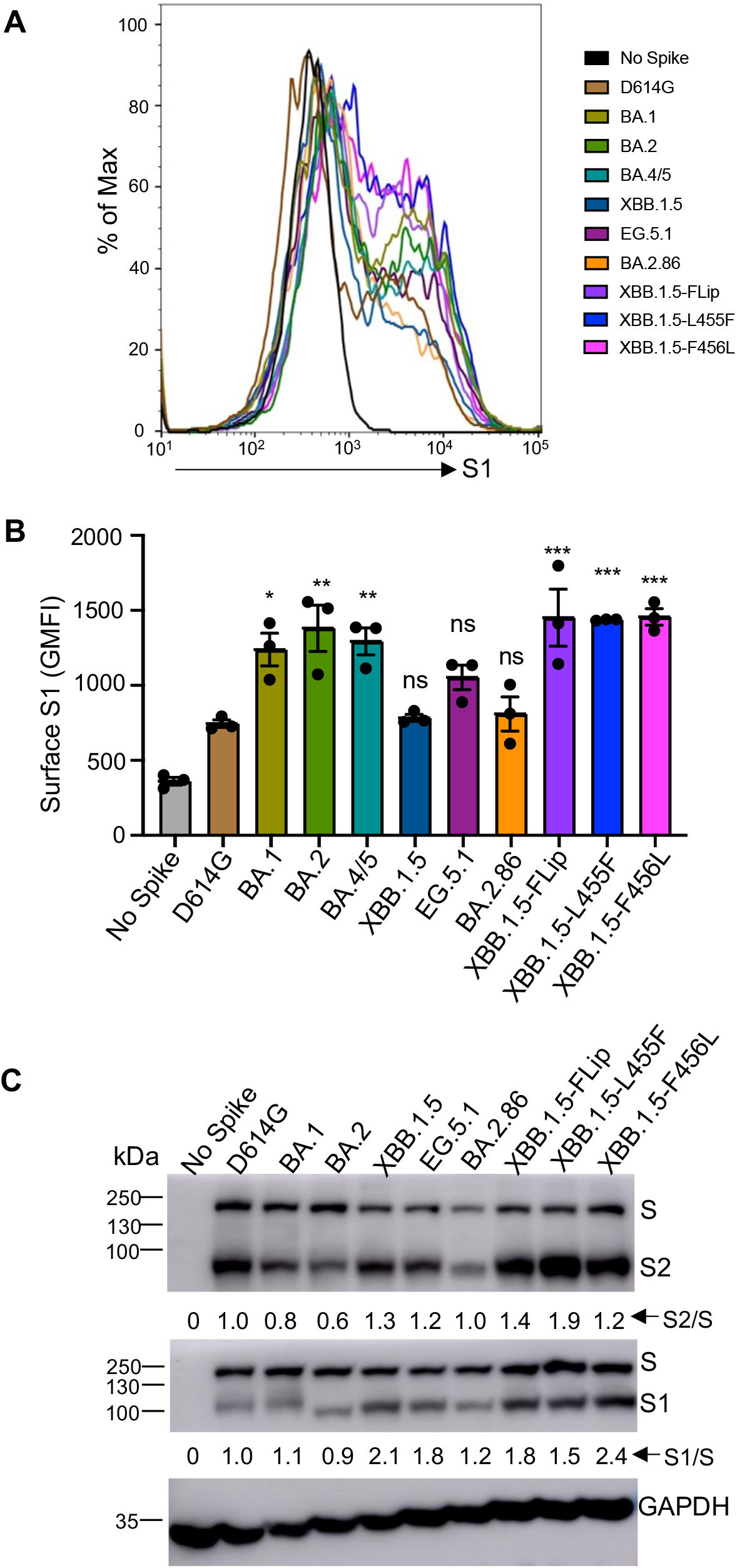
Cell surface expression and processing of Omicron BA.2.86 and Flip Spike proteins. (**A-B**) Cell surface expression of the indicated variant Spike proteins. HEK293T cells used for production of pseudotyped lentiviral vectors carrying each variant Spike proteins (Figures 1-3) were stained with anti-SARS-CoV-2 S1 antibody. Representative histogram of anti-S1 signals in the cells (**A**) and geometric mean fluorescence intensities (**B**) of each subvariant from three biological replicates are shown. (**C**) Spike expression and processing in viral producer cell lysates. HEK293T cells, which were used to produce lentiviral pseudotypes, were lysed and probed with anti-S1, anti-S2 and anti-GAPDH antibodies, respectively. Spike processing was quantified by NIH ImageJ and set to a S1/S or S2/S ratio and normalized the ratios of each Omicron subvariant to that of D614G. Dots represent three biological replicates. Bars in (**B**) represent means ± standard error. Significance relative to D614G was made using a one-way ANOVA with Bonferroni post-test. P values are displayed as ns p > 0.05, *p < 0.05, **p < 0.01, and ***p < 0.001.

### Molecular modeling revealed how mutations in BA.2.86 compromise S309 antibody neutralization

We performed homology modeling to understand the possible molecular and structural basis by which BA.2.86 exhibits distinct viral infectivity and evades S309 neutralization. **Fig. 6A** shows a model of the BA.2.86 Spike trimer, highlighting mutations that differ from the ancestral BA.2 variant. S309, classified as a class III monoclonal antibody, targets the lateral segment of the Receptor Binding Domain (RBD) within the Spike protein, especially residues 330-441. Among these residues, positions 339 and 356 are pivotal components of the S309- binding epitope. The replacement of the native Glycine 339 residue with either Aspartic acid (present in BA.2 and BA.4/5) or Histidine (present in BA.2.86, XBB.1.5, and EG.5.1) creates steric hindrance effects that interfere binding with residues Y100 and L110 of the S309 antibody (**Fig. 6B**). Simultaneously, the K356T mutation, which is also present in BA.2.86, disrupts the salt-bridge interaction established with E108 of S309 (**Fig. 6B**). Together, these dual mutations diminish the neutralization efficacy of the BA.2.86 variant by the S309 antibody.

**Figure 6:**
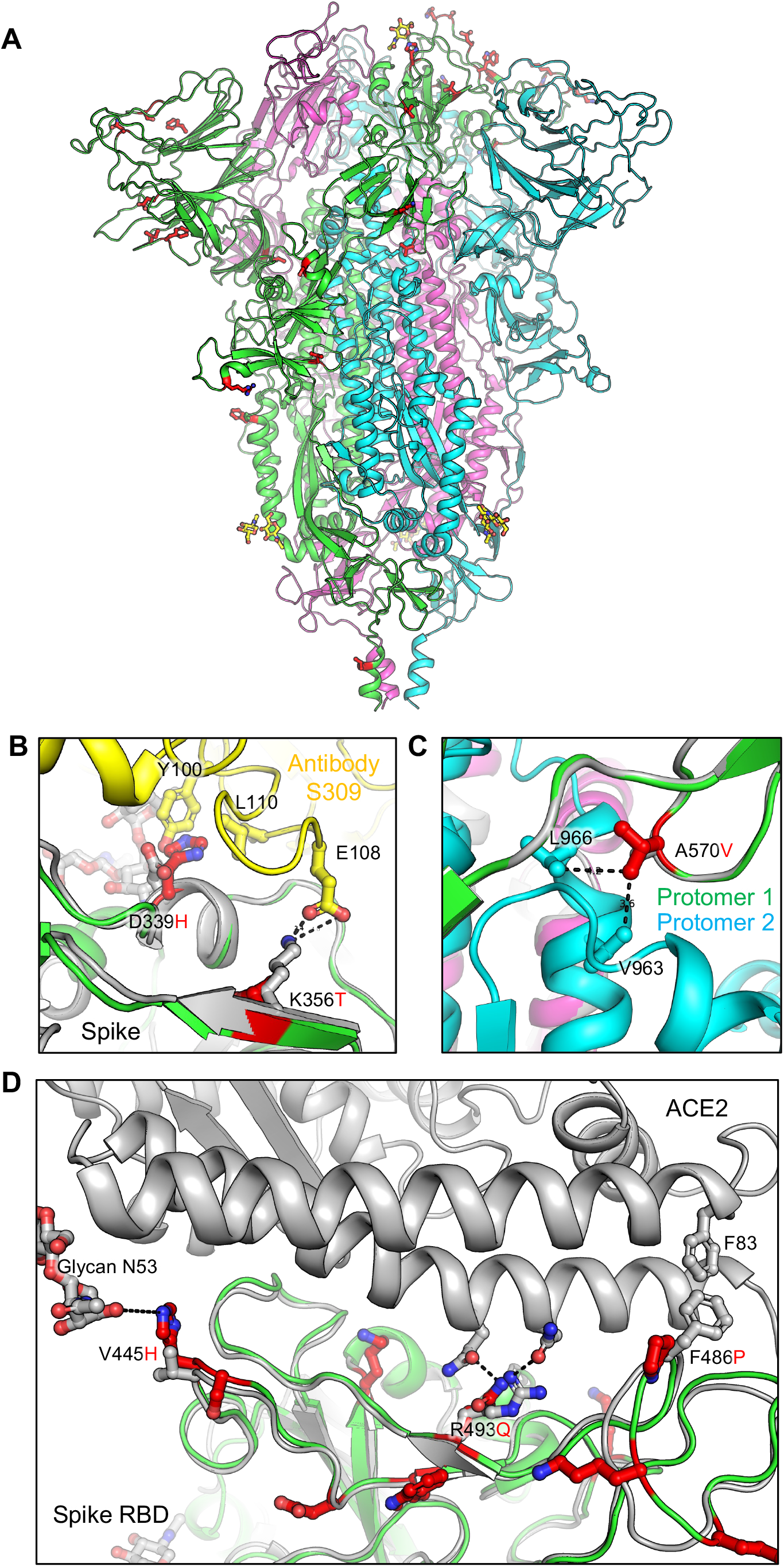
Homology modeling of key mutations in BA.2.86. (**A**) A homology model of the BA.2.86 Spike trimer is presented, highlighting mutations that differ from the BA.2 variant as red sticks on the green protomer. (**B**) The substitution of the wildtype G339 residue with either D or H introduces steric hindrance to residues Y100 and L110 of antibody S309. Simultaneously, the K356T mutation disrupts the salt-bridge interaction with E108 of S309. These mutations collectively impair the recognition of the Spike protein by antibody S309. (**C)** The A570V mutation in BA.2.86 Spike enhances hydrophobic interactions between protomers, thereby increasing trimer stability. (**D**) Focusing on the RBM region, the V445H and R493Q mutations may enhance receptor binding by introducing hydrogen bonds between the Spike protein and the ACE2 receptor. Conversely, the F486P mutation weakens receptor binding by losing the hydrophobic interaction with F83 of ACE2.

### Antigenic mapping shows distinct antigenicity of BA.2.86 from FLip and other XBB variants

We next analyzed the extent to which antigenicity of the different Spikes varies using antigenic mapping analysis on our three cohorts of neutralization data shown in **Fig. 2**. This analysis is adopted from a study by Smith and colleagues investigating the antigenicity of different influenza hemagglutinin proteins based on agglutination neutralization assays^43^. To construct the maps, we used the Racmacs program, which performs multidimensional scaling on log2 transformed neutralization assay results. These calculations were used to construct maps that plot points for individual antigens and antibodies in Euclidean space^43^. Therefore, the spaces between these points are directly related to fold changes in neutralization titers, allowing for a visual representation of the antigenic distance between variant Spikes in our assay. Note that the plots are constructed in units of “antigenic distance units” (AU) where 1 AU represents a 2-fold change in neutralizing antibody titer^13,43^. For all cohorts, D614G, BA.1, and BA.2 Spikes consistently cluster together with BA.4/5 nearby (**Fig. 7A-C**); XBB lineage variants cluster farther away, averaging about 5-7 AU away from D614G which translates to 32∼128- fold changes in neutralization titers (**Fig. 7A-C**, **Fig. 2**). The antigenic distance between variants decreases from the 3-dose vaccinated plot to the bivalent vaccinated plot (**Fig 7A-B**), suggesting that the dose of bivalent vaccine broadens the immune response against Omicron subvariants. For all cohorts, BA.2.86 is antigenically more similar to D614G, with antigenic distances of 3.5∼5.5 AU from D614G, whereas the FLip variant is more antigenically distinct from D614G and early Omicron subvariants with antigenic distances of 6∼7 AU from D614G (**Fig 7A-C**). Overall, this analysis suggests that BA.2.86 is more antigenically similar to early Omicron subvariants BA.1, BA.2, and BA.4/5 and antigenically distinct from the FLip variant.

**Figure 7:**
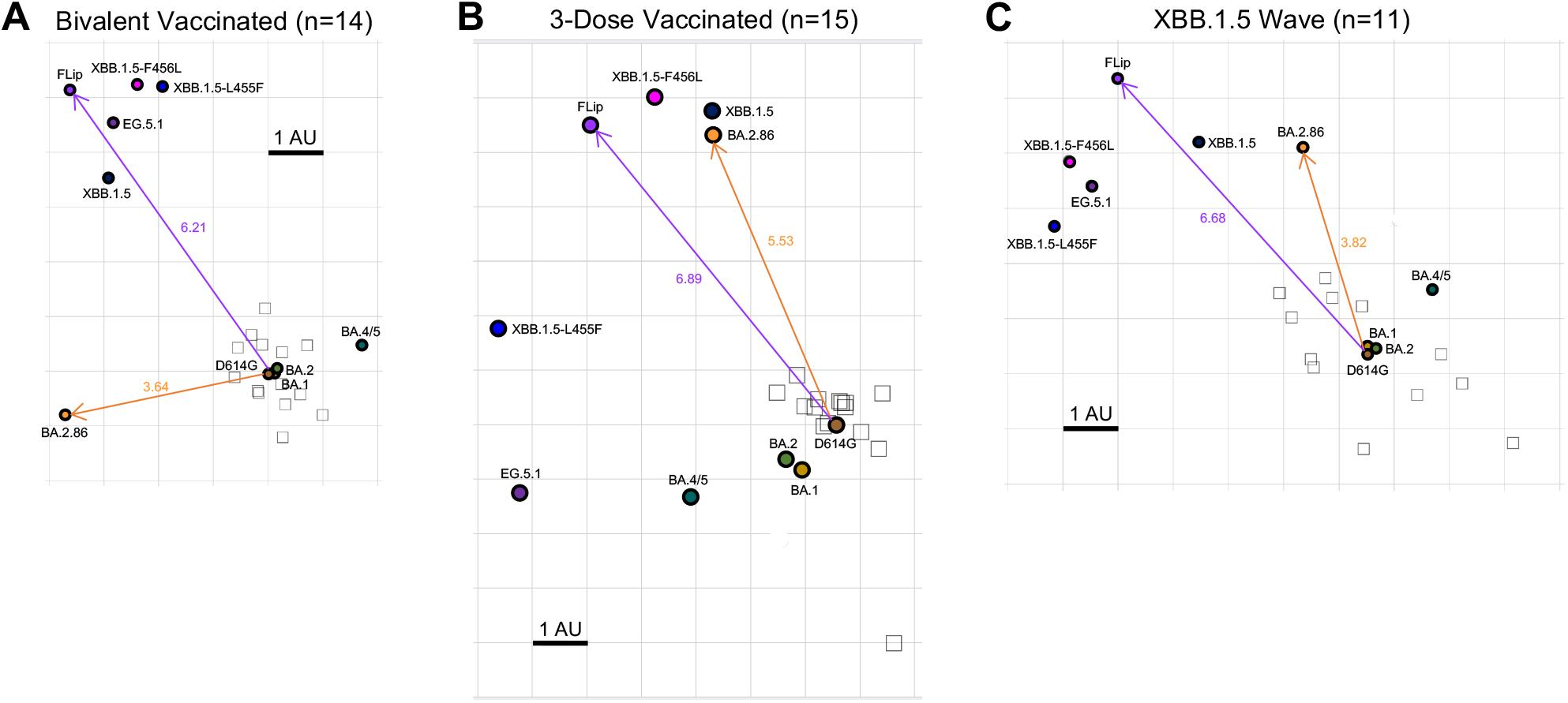
Antigenic mapping of neutralization titers for bivalent vaccinated, monovalent vaccinated, and XBB.1.5-wave infected cohorts, derived from the NT_50_ in. Figure 2. Antigenic maps for neutralization titers from **(A)** the bivalent vaccinated, **(B)** the monovalent vaccinated, and **(C)** the XBB.1.5-wave infected cohorts were made using the Racmacs program (1.1.35) (see Methods). Squares represent the individual sera sample and circles represent the variants. One square on the grid represents one antigenic unit squared.

## Discussion

The ongoing evolution of SARS-CoV-2 has presented a constant challenge for the development of effective COVID-19 vaccines. Here, we characterized the neutralization of two new variants, BA.2.86 and FLip, by bivalent-vaccinated, monovalent-vaccinated, or XBB.1.5-infected sera, as well as by the monoclonal antibody S309. We showed that, while 3 doses of monovalent vaccine remain ineffective against BA.2.86, FLip and other XBB subvariants, the bivalent-vaccinated sera can efficiently neutralize BA.2.86, with nAb titers actually higher than that of XBB.1.5, EG.5.1, and FLip. A similar trend was observed for the XBB.1.5-wave cohort – despite generally low titers, especially in those who had been vaccinated with bivalent vaccines. These results are somewhat surprising, given that BA.2.86 has >30 mutations relative to XBB variants; however, our data are consistent with those of other groups (preprints). Together, these results support the conclusion that BA.2.86 is not as immune evasive as the XBB variants, especially FLip and EG.5.1, which may also explain, in part, why BA.2.86 has not risen to as much dominance in circulation as the original Omicron did. While BA.2.86 appears to exhibit quite distinct antigenicity (**Fig. 7**)^32^, it is closer to the early Omicron BA.1, BA.2 and BA.4/5 in contrast to XBB variants, especially FLip. Interestingly, sera from individuals vaccinated with the new Moderna monovalent XBB.1.5 mRNA vaccine have shown robust and comparable efficacy against BA.2.86 and XBB including FLip^44^.

Vaccination is critical for protection against COVID-19, but monoclonal antibodies also play an important role. Unfortunately, many monoclonal antibodies have lost the ability to neutralize Spike upon emergence of new Omicron variants. S309, a class III antibody, however, has largely maintained efficacy against Omicron Spike lineages^16,40,41^, with notable exceptions of BA.2.75.2, CH.1.1, and CA.3.1 as shown in our previous study^16^, likely due to mutations at residues 346 and 339 of the Spike. In this work, we found that S309 is unable to neutralize BA.2.86, which also has the Spike mutation D339H located within the epitope binding region for S309, as shown in our model (**Fig. 6B**). Moreover, a second mutation K356T abolish the important hydrophilic interaction to this antibody. These dual mutations significantly impair the neutralization efficiency of antibody S309. Further studies are needed to confirm the role of the dual mutation in facilitating BA.2.86 evasion of the neutralization by S309 as well as possible roles of other Spike mutations in BA2.86.

Interestingly, BA.2.86 presents distinct biology from BA.2 and XBB variants. We have previously shown that the original BA.1/BA.2 Omicron Spike has low infectivity in CaLu-3 cells, decreased fusogenicity in 293T- ACE2 cells, and impaired Spike processing in virus producer cells^5,11^. Here we find that BA.2.86 displays decreased infectivity in 293T-ACE2 cells, not only compared to the ancestral BA.2/BA.1 but also relative to more recent XBB variants, including XBB.1.5, EG.5.1 and FLip. Moreover, the fusion activity of BA.2.86 is also low in 293T-ACE2 cells, consistent with the relatively low efficiency of Spike processing as well as surface expression. Strikingly, in CaLu-3 cells, BA.2.86 exhibits a higher infectivity as well as enhanced cell-cell fusion compared to the ancestral BA.2 and some XBB variants. These results suggest that the Spike protein of BA.2.86 may be more conformational stable compared to the parental BA.2 and XBB variants, especially FLip and EG.5.1. Indeed, molecular modeling shows that the A570V mutation enhances hydrophobic interactions between protomers, thereby potentially increasing trimer stability (**Fig 6C**). However, the exact mechanisms underlying the distinct fusogenicity and/or stability of BA.2.86 will be investigated in future studies.

The increased infectivity of BA.2.86 in CaLu-3 cells is somewhat alarming. CaLu-3 represents a biologically relevant cell line that is derived from human lung epithelia type II pneumocytes and is known to express endogenous levels of ACE2 and host co-factor TMPRSS2 — the latter is critical for the respiratory tract tropism for SARS-CoV-2^4,45–47^. It has been established that CaLu-3 cells are almost exclusively infected through the TMPRSS2-reliant plasma membrane fusion pathway, while the endosomal pathway is used in 293T-ACE2 cells due to the lack of TMPRSS2. Furthermore, comparisons between the Delta and Omicron variants demonstrated that Omicron^5,46,48^ associates with increased transmissibility^2,46^, but decreased pathogenicity versus Delta^2–4^. Our data shown here suggest that BA.2.86 may have an increased tendency of using the plasma membrane route of entry, as opposed to the endosomal route of entry. Our molecular modeling suggests that mutations present in BA.2.86 and XBB variants can alter the Spike binding to ACE2 receptor, therefore impacting membrane fusion and entry of different target cells. For example, the V445H and R493Q mutations may enhance ACE2 binding by introducing hydrogen bonds between the Spike protein of BA.2.86/XBB.1.5 and the ACE2 receptor. Conversely, the F486P mutation present in XBB.1.5 weakens receptor binding by losing the hydrophobic interaction with F83 of ACE2 (**Fig. 6D**). Whether or not BA.2.86 will have an increased lung tropism, thus enhanced pathogenesis compared to other Omicron variants, is unknown and needs to be carefully examined.

## Acknowledgements

We wish to thank the Clinical Research Center/Center for Clinical Research Management of The Ohio State University Wexner Medical Center and The Ohio State University College of Medicine in Columbus, Ohio, specifically J. Brandon Massengill, Francesca Madiai, Dina McGowan, Breona Edwards, Evan Long, and Trina Wemlinger, for collection and processing of samples. We also thank Tongqing Zhou at NIH for providing the S309 monoclonal antibody. In addition, we thank Sarah Karow, Madison So, Preston So, Daniela Farkas, and Finny Johns in the clinical trials team of The Ohio State University for sample collection and other supports. In addition, we thank Moemen Eltobgy for assistance in sample processing. We specially thank Ashish R. Panchal, Soledad Fernandez, Mirela Anghelina, and Patrick Stevens for their assistance in providing the sample information of the first responders and their household contacts. We thank Peng Ru and Lauren Masters for sequencing and Xiaokang Pan for bioinformatic analysis. S.-L.L., D. J., R.J.G., L.J.S. and E.M.O. were supported by the National Cancer Institute of the NIH under award no. U54CA260582. The content is solely the responsibility of the authors and does not necessarily represent the official views of the National Institutes of Health. This work was also supported by a fund provided by an anonymous private donor to OSU. P.Q. was supported by a Glenn Barber Fellowship from the Ohio State University College of Veterinary Medicine. K.X. was supported by The Ohio State University Comprehensive Cancer Center, and a Path to K grant through the Ohio State University Center for Clinical & Translational Science. J.S.B. was supported by Award Number Grant UL1TR002733 and KL2TR002734 from the National Center for Advancing Translational Sciences. R.J.G. was additionally supported by the Robert J. Anthony Fund for Cardiovascular Research and the JB Cardiovascular Research Fund, and L.J.S. was partially supported by NIH R01 HD095881.

## Author contributions

S.-L.L. conceived and directed the project. R.J.G led the clinical study/experimental design and implementation. P.Q. performed most of the experiments, J.N.F performed antigenic mapping, N.G. performed mutagenesis and sequencing of new variants. P.Q and J.N.F. performed data processing and analyses. D.J. led SARS-CoV-2 variant genotyping and DNA sequencing analyses. C.C., J.S.B, J.C.H., R.M. and R.J.G. provided clinical samples and related information. K.X. performed molecular modeling and participated in discussion. P.Q., J.N.F., and S.-L.L. wrote the paper. Y.-M.Z, L.J.S., and E.M.O. provided insightful discussion and revision of the manuscript.

## Study cohorts, Materials, and Methods

### Vaccinated and convalescent cohorts

This study included three different groups of human sera that were tested for neutralizing antibody titers against the selected panel of SARS-CoV-2 variants. The first cohort were HCWs working at the Ohio State Wexner Medical Center that received 3 homologous doses of monovalent mRNA vaccine (n=15). Samples were collected under the approved IRB protocols 2020H0228, 2020H0527, and 2017H0292. Of the 15 total individuals, 3 received the Moderna mRNA-1273 vaccine and 12 received Pfizer BioNTech BNT162b2 vaccine. Sera samples were collected between 14-86 days after administration of the third vaccine dose. Individuals ranged from 26-61 years old (median 33), 10 males and 5 females were included.

The second cohort were HCWs working at the Ohio State Wexner Medical Center that received at least 2 doses of monovalent vaccine and 1 dose of bivalent vaccine (n=14). Samples were collected under the approved IRB protocols 2020H0228, 2020H0527, and 2017H0292. 12 individuals received 3 doses of homologous monovalent mRNA vaccine (Pfizer or Moderna) with an additional 1 dose of bivalent vaccine (Pfizer). 1 person received 4 doses of monovalent vaccine (Pfizer) and a bivalent booster (Pfizer) and the last person received 2 doses of monovalent vaccine (Pfizer) and 1 bivalent booster (Pfizer). Sera samples were collected between 23-108 days post bivalent dose administration. Individuals ranged from 25-48 years old, 8 males and 6 females were included.

The last cohort included were first responders that were infected during the XBB.1.5 wave of infection (n=8) and hospitalized patients (n=3) in Columbus, Ohio (February 2023-Late August 2023) (total n=11). Samples were collected under IRB protocols 2020H0527, 2020H0531, 2020H0240, and 2020H0175. Nasal swabs were used to confirm COVID-19 positive status and were also used for sequencing to determine the infecting variant. Eight of the samples were confirmed to be XBB.1.5 using COVID-Seq Artic v4 sequencing and typed with Dragen COVID Lineage with Pangolin plug-in (Illumina). 3 sample did not have conclusive sequencing but largely matched with XBB.1 lineage and aligned with the timing of the XBB.1.5 wave. Additional 8 individuals were vaccinated and 3 were completely unvaccinated. Vaccinated people included 2 that received two doses of monovalent vaccine (1 Moderna, 1 Janseen), 3 people that received 3 doses of monovalent vaccine (1 Moderna, 2 Pfizer), 1 that received 3 doses of Moderna vaccine and 1 dose of Moderna bivalent booster, 2 person that had 4 doses of Moderna monovalent vaccine 1 dose of Pfizer bivalent booster. Individuals ranged from 36-75 years old (median 53), 7 males and 4 females were included.

### Cell lines

The cell lines used included human embryonic kidney 293T cells (ATCC CRL-11268, RRID: CVCL_1926), HEK293T cells expressing human ACE2 (BEI NR-52511, RRID: CVCL_A7UK), and human lung adenocarcinoma cell line CaLu-3 (RRID: CVCL_0609). HEK293T cells were cultured in DMEM (Gibco, 11965-092) plus 10% FBS (Sigma, F1051) and 0.5% penicillin-streptomycin (HyClone, SV30010). CaLu-3 cells were maintained in EMEM (ATCC, 30-2003) supplemented the same way. To split, cells were washed in phosphate buffered saline (Sigma, D5652-10X1L) then incubated in 0.05% trypsin + 0.53 mM EDTA (Corning, 25-052-CI) until complete detachment.

### Plasmids

Plasmids used in this study include the individual spike plasmids engineered in the pcDNA3.1 backbone, the pNL4-3-inGluc lentiviral vector, and eGFP. Spike plasmids include FLAG tags and were either engineered by GenScript Biotech (Piscataway, NJ) through restriction enzyme cloning (D614G, BA.1, BA.2, BA.4/5, BA.2.86) or generated by our lab through site-directed mutagenesis (XBB.1.5, EG.5.1, FLip, XBB.1.5-L455F, XBB.1.5- F456L). The pNL4-3 vector is an HIV-1 vector with an Env deletion and a *Gaussia* luciferase reporter interrupted by an intron as described in a previous study^5^.

### Pseudotyped lentivirus production and infectivity

Pseudotyped lentiviral vectors were produced by cotransfecting 293T cells with pNL4-3-inGluc vector and spike in a 2:1 ratio. Polyethyleneimine transfection was used (Transporter 5 Transfection Reagent, Polysciences). Pseudotyped vectors were collected by taking the media off producer cells at 48 and 72 hours post-transfection. The collected vectors were then used to infect either HEK293T-ACE2 or CaLu-3 cells. Infectivity was measured through relative luminescence readouts by taking infected cell media and combining it with an equal volume of luciferase substrate (0.1 M Tris pH 7.4, 0.3 M sodium ascorbate, 10 µM coelenterazine). Readings were measured with a BioTek Cytation plate reader.

### Virus neutralization assay

Sera samples were diluted 1:40 then serially diluted for final concentrations 1:40, 1:160, 1:640, 1:2560, 1:10240, and no serum as a control. mAb S309 was diluted 12 μg/mL then diluted 4-fold for final concentrations 12, 3, 0.75, 0.1875, 0.046875 μg/ml, no antibody control. The collected pseudotyped virus was titered as described previously and diluted to normalize any variation in titer. 100 uL of normalized virus was mixed with the sera samples and incubated for 1 hour and 37C. After the incubation, this mixture was used to infect 293T- ACE2 cells. Luminescence readouts were collected 48 and 72 hours post-infection and used to calculate neutralization titers at 50% (NT_50_). NT_50_ values were calculated using least-squares fit non-linear regression with normalized response (no serum control) using GraphPad Prism v9 (San Diego, CA).

### Cell-cell fusion

To assess fusogenicity of the spikes, 293T cells were cotransfected with eGFP and Spike of interest. Next day, the effector 293T cells were digested by trypsin and cocultured with digested 293T-ACE2 or CaLu-3 cells. Spike expressed on the membrane of cells was allowed to interact with ACE2 on neighboring cells and trigger the cell-cell fusion over the course of 24 hours. Cell-cell fusion were imaged using a Leica DMi8 microscope and the Leica X Applications Suite software was used to outline the edges of syncytia based on the GFP signal and calculate the area of the fused cell bodies. Three images from duplicate wells were randomly taken. Scale bars represent 150 µM and one representative image was selected for presentation.

### Spike protein surface expression

After collecting virus from the 293T cells used to produce the lentiviral vectors, the producer cells were washed in PBS and detached using PBS + 5mM EDTA. A portion of these cells were taken and fixed using 3.7% formaldehyde for 10 minutes at room temperature. Cells were stained with polyclonal anti-S1 antibody (Sino Biological, 40591-T62; RRID: AB_2893171) for 1.5 hours and washed three times with PBS+2% FBS. The secondary stain used was anti-Rabbit-IgG-FITC (Sigma, F9887, RRID: AB_259816). Cells were then washed another 3 times then flow cytometry was performed using a LifeTechnologies Attune NxT flow cytometer. Data analysis was conducted using FlowJo v10.9.1 (Ashland, OR).

### Spike protein processing

The remaining virus producer cells leftover after taking cells for flow cyometry were lysed using RIPA buffer supplemented with protease inhibitor cocktails (RIPA: 50 mM Tris pH 7.5, 150 mM NaCl, 1 mM EDTA, Nonidet P-40, 0.1% SDS, PI+PMSF: Sigma, P8340) for 40 minutes on ice. Lysate was then harvested and used for western blotting. Samples were separated using a 10% acrylamide SDS-PAGE gel and transferred to a PVDF membrane. Blots were then incubated with polyclonal anti-S2 antibody (Sino Biological, 40590; RRID:AB_2857932), anti-S1 antibody (Sino Biological, 40591-T62; RRID: AB_2893171), and anti-GAPDH as a loading control (Santa Cruz, Cat# sc-47724, RRID: AB_627678). Secondary antibodies used were anti-Rabbit-IgG-FITC (Sigma, A9169; RRID:AB_258434) and anti-Mouse-IgG-FITC (Sigma, Cat# A5278, RRID: AB_258232). Blots were imaged using Immobolin Crescendo Western HRP substrate (Millipore, WBLUR0500) and exposed on a GE Amersham Imager 600. Quantification of band intensity was determined using ImageJ (NIH, Bethesda, MD).

### Homology modeling

Structural modeling of the BA.2.86 Spike protein was used to explore how it interacts with both the ACE2 receptor and neutralizing antibodies. This was performed by the SWISS-MODEL server with published structures from X-ray crystallography or cryo-EM studies (PDB: 7XOC, 7XCK, 7R6W, 7XIX, 7XIW) as templates. Key mutations affecting the potential interactions were examined and presented visually with PyMOL.

### Antigenic mapping

Antigenic mapping was performed using the Racmacs program (v1.1.35) (https://github.com/acorg/Racmacs/tree/master) in R (Vienna, Austria). The program works by converting raw neutralization titers into log2 transformed values and using them to generate a distance table for the individual antigen and sera values. The program then performs multidimensional scaling based on the table to generate a map where each antigen and sera sample is represented by a single point in Euclidean space and distance between them directly correlates to antigenic differences. 1 antigenic distance unit (AU) is equivalent to a 2-fold change in neutralizing antibody titer^13,43^. Optimization settings for mapping were kept on default (2 dimensions, 500 optimizations, minimum column basis “none”). Maps were saved from the “view(map)” function and labels were added using Microsoft Office PowerPoint. The length of arrows drawn within PowerPoint between antigen points were used to calculate the distance between points. These distances were normalized using the scale bar for “1 AU.”

### Quantification and statistical analysis

All statistical analyses that were described in the figure legends were conducted using GraphPad Prism 9. NT_50_ values were calculated by least-squares fit non-linear regression. Error bars in **(Fig. 1B-C**, **Fig. 3A**, **Fig. 4B**, **Fig. 4D**, **Fig.5B, and Fig. S1A-B)** represent means ± standard error. Error bars in **(Fig. 2A**, **Fig. 2C**, **Fig. 2E** and **Fig. S1A-B)** represent geometric means with 95% confidence intervals. Statistical significance was analyzed using log10 transformed NT_50_ values to better approximate normality (**Fig. 2A**, **Fig. 2C**, **Fig. 2E** and **Fig. S1A-B**), and multiple groups comparisons were made using a one-way ANOVA with Bonferroni post-test. Cell-cell fusion were quantified using the Leica X Applications Suite software (**Fig. 4B** and **Fig. 4D**). S processing was quantified by NIH ImageJ (**Fig. 5C**).

## References

1. Gangavarapu, K., Latif, A., Mullen, J., Alkuzweny, M., Hufbauer, E., Tsueng, G., Haag, E., Zeller, M., Aceves, C., and Zaiets, K. (2023). SARS-CoV-2 (hCoV-19) Mutation Reports.

2. Yuan, S., Ye, Z.-W., Liang, R., Tang, K., Zhang, A.J., Lu, G., Ong, C.P., Man Poon, V.K., Chan, C.C.-S., Mok, B.W.-Y., et al. (2022). Pathogenicity, transmissibility, and fitness of SARS-CoV-2 Omicron in Syrian hamsters. Science 377, 428–433. doi:10.1126/science.abn8939.

3. Suzuki, R., Yamasoba, D., Kimura, I., Wang, L., Kishimoto, M., Ito, J., Morioka, Y., Nao, N., Nasser, H., Uriu, K., et al. (2022). Attenuated fusogenicity and pathogenicity of SARS-CoV-2 Omicron variant. Nature 603, 700–705. 10.1038/s41586-022-04462-1.

4. Shuai, H., Chan, J.F., Hu, B., Chai, Y., Yuen, T.T., Yin, F., Huang, X., Yoon, C., Hu, J.C., Liu, H., et al. (2022). Attenuated replication and pathogenicity of SARS-CoV-2 B.1.1.529 Omicron. Nature 603, 693–699. 10.1038/s41586-022-04442-5.

5. Zeng, C., Evans, J.P., Qu, P., Faraone, J., Zheng, Y.M., Carlin, C., Bednash, J.S., Zhou, T., Lozanski, G., Mallampalli, R., et al. (2021). Neutralization and Stability of SARS-CoV-2 Omicron Variant. bioRxiv. 10.1101/2021.12.16.472934.

6. Xia, H., Zou, J., Kurhade, C., Cai, H., Yang, Q., Cutler, M., Cooper, D., Muik, A., Jansen, K.U., Xie, X., et al. (2022). Neutralization and durability of 2 or 3 doses of the BNT162b2 vaccine against Omicron SARS-CoV-2. Cell Host Microbe 30, 485–488.e483. 10.1016/j.chom.2022.02.015.

7. Wang, X., Zhao, X., Song, J., Wu, J., Zhu, Y., Li, M., Cui, Y., Chen, Y., Yang, L., Liu, J., et al. (2022). Homologous or heterologous booster of inactivated vaccine reduces SARS-CoV-2 Omicron variant escape from neutralizing antibodies. Emerg Microbes Infect 11, 477–481. 10.1080/22221751.2022.2030200.

8. Schmidt, F., Muecksch, F., Weisblum, Y., Da Silva, J., Bednarski, E., Cho, A., Wang, Z., Gaebler, C., Caskey, M., Nussenzweig, M.C., et al. (2022). Plasma Neutralization of the SARS-CoV-2 Omicron Variant. N Engl J Med 386, 599–601. 10.1056/NEJMc2119641.

9. Planas, D., Saunders, N., Maes, P., Guivel-Benhassine, F., Planchais, C., Buchrieser, J., Bolland, W.H., Porrot, F., Staropoli, I., Lemoine, F., et al. (2022). Considerable escape of SARS-CoV-2 Omicron to antibody neutralization. Nature 602, 671–675. 10.1038/s41586-021-04389-z.

10. Pérez-Then, E., Lucas, C., Monteiro, V.S., Miric, M., Brache, V., Cochon, L., Vogels, C.B.F., Malik, A.A., De la Cruz, E., Jorge, A., et al. (2022). Neutralizing antibodies against the SARS-CoV-2 Delta and Omicron variants following heterologous CoronaVac plus BNT162b2 booster vaccination. Nature medicine 28, 481–485. 10.1038/s41591-022-01705-6.

11. Evans, J.P., Zeng, C., Qu, P., Faraone, J., Zheng, Y.-M., Carlin, C., Bednash, J.S., Zhou, T., Lozanski, G., Mallampalli, R., et al. (2022). Neutralization of SARS-CoV-2 Omicron sub-lineages BA.1, BA.1.1, and BA.2. Cell host & microbe 30, 1093–1102.e1093. 10.1016/j.chom.2022.04.014.

12. Zou, J., Kurhade, C., Patel, S., Kitchin, N., Tompkins, K., Cutler, M., Cooper, D., Yang, Q., Cai, H., Muik, A., et al. (2023). Neutralization of BA.4-BA.5, BA.4.6, BA.2.75.2, BQ.1.1, and XBB.1 with Bivalent Vaccine. N Engl J Med 388, 854–857. 10.1056/NEJMc2214916.

13. Wang, Q., Iketani, S., Li, Z., Liu, L., Guo, Y., Huang, Y., Bowen, A.D., Liu, M., Wang, M., Yu, J., et al. (2023). Alarming antibody evasion properties of rising SARS-CoV-2 BQ and XBB subvariants. Cell 186, 279–286.e278. 10.1016/j.cell.2022.12.018.

14. Wang, Q., Guo, Y., Zhang, R.M., Ho, J., Mohri, H., Valdez, R., Manthei, D.M., Gordon, A., Liu, L., and Ho, D.D. (2023). Antibody Neutralization of Emerging SARS-CoV-2: EG.5.1 and XBC.1.6. bioRxiv, 2023.2008.2021.553968. 10.1101/2023.08.21.553968.

15. Uraki, R., Ito, M., Furusawa, Y., Yamayoshi, S., Iwatsuki-Horimoto, K., Adachi, E., Saito, M., Koga, M., Tsutsumi, T., Yamamoto, S., et al. (2023). Humoral immune evasion of the omicron subvariants BQ.1.1 and XBB. Lancet Infect Dis 23, 30–32. 10.1016/s1473-3099(22)00816-7.

16. Qu, P., Faraone, J.N., Evans, J.P., Zheng, Y.-M., Carlin, C., Anghelina, M., Stevens, P., Fernandez, S., Jones, D., Panchal, A.R., et al. (2023). Enhanced evasion of neutralizing antibody response by Omicron XBB.1.5, CH.1.1, and CA.3.1 variants. Cell Reports 42, 112443.

17. Miller, J., Hachmann, N.P., Collier, A.Y., Lasrado, N., Mazurek, C.R., Patio, R.C., Powers, O., Surve, N., Theiler, J., Korber, B., and Barouch, D.H. (2023). Substantial Neutralization Escape by SARS-CoV-2 Omicron Variants BQ.1.1 and XBB.1. N Engl J Med 388, 662–664. 10.1056/NEJMc2214314.

18. Kurhade, C., Zou, J., Xia, H., Liu, M., Chang, H.C., Ren, P., Xie, X., and Shi, P.Y. (2022). Low neutralization of SARS-CoV-2 Omicron BA.2.75.2, BQ.1.1 and XBB.1 by parental mRNA vaccine or a BA.5 bivalent booster. Nat Med. 10.1038/s41591-022-02162-x.

19. Imai, M., Ito, M., Kiso, M., Yamayoshi, S., Uraki, R., Fukushi, S., Watanabe, S., Suzuki, T., Maeda, K., Sakai-Tagawa, Y., et al. (2023). Efficacy of Antiviral Agents against Omicron Subvariants BQ.1.1 and XBB. N Engl J Med 388, 89–91. 10.1056/NEJMc2214302.

20. Faraone, J.N., Qu, P., Evans, J.P., Zheng, Y.-M., Carlin, C., Anghelina, M., Stevens, P., Fernandez, S., Jones, D., Lozanski, G., et al. (2023). Neutralization escape of Omicron XBB, BR.2, and BA.2.3.20 subvariants. Cell Reports Medicine 4, 101049.

21. Davis-Gardner, M.E., Lai, L., Wali, B., Samaha, H., Solis, D., Lee, M., Porter-Morrison, A., Hentenaar, I.T., Yamamoto, F., Godbole, S., et al. (2022). Neutralization against BA.2.75.2, BQ.1.1, and XBB from mRNA Bivalent Booster. New England Journal of Medicine 388, 183–185. 10.1056/NEJMc2214293.

22. Pfizer and BioNTech Submit Applications to U.S. FDA for Omicron XBB.1.5-Adapted Monovalent COVID-19 Vaccine. (2023).

23. MODERNA FILES FOR FDA AUTHORIZATION OF ITS UPDATED COVID-19 VACCINE. (2023).

24. Recommendation for the 2023-2024 Formula of COVID-19 vaccines in the U.S. (2023).

25. Risk assessment for SARS-CoV-2 variant V-23AUG-01 (or BA.2.86). (2023).

26. Risk Assessment Summary for SARS CoV-2 Sublineage BA.2.86. (2023).

27. Callaway, E. (2023). Why a highly mutated coronavirus variant has scientists on alert. News Explainer.

28. Topol, E. (2023). A Quick Update on the BA.2.86 Variant. Ground Truths. Substack.

29. Schnirring, L. (2023). A few more BA.2.86 COVID-19 detections noted in human samples, wastewater. CIDRAP, 8/28/2023.

30. Schnirring, L. (2023). WHO adds BA.2.86 to SARS-CoV-2 variant monitoring list. CIDRAP, 8/17/2023.

31. Looi, M.-K. (2023). Covid-19: Scientists sound alarm over new BA.2.86 “Pirola” variant. BMJ 382, p1964. 10.1136/bmj.p1964.

32. Bloom, J. (2023). Phenotypic assessment of spike mutations in BA.2.86 (new highly mutated BA.2 variant).

33. SARS-CoV-2 variant surveillance and assessment: technical briefing 53. (2023).

34. Update on SARS CoV-2 Variant BA.2.86. (2023).

35. CDC COVID Data Tracker: Variant Proportions. (2023).

36. Faraone, J.N., Qu, P., Goodarzi, N., Zheng, Y.-M., Carlin, C., Saif, L.J., Oltz, E.M., Xu, K., Jones, D., Gumina, R.J., and Liu, S.-L. (2023). Immune Evasion and Membrane Fusion of SARS-CoV-2 XBB Subvariants EG.5.1 and XBB.2.3. bioRxiv, 2023.2008.2030.555188. 10.1101/2023.08.30.555188.

37. Yang, S., Yu, Y., Jian, F., Song, W., Yisimayi, A., Chen, X., Xu, Y., Wang, P., Wang, J., Yu, L., et al. (2023). Antigenicity and infectivity characterization of SARS-CoV-2 BA.2.86. bioRxiv, 2023.2009.2001.555815. 10.1101/2023.09.01.555815.

38. Sheward, D.J., Yang, Y., Westerberg, M., Öling, S., Muschiol, S., Sato, K., Peacock, T.P., Hedestam, G.B.K., Albert, J., and Murrell, B. (2023). Sensitivity of BA.2.86 to prevailing neutralising antibody responses. bioRxiv, 2023.2009.2002.556033. 10.1101/2023.09.02.556033.

39. San Filippo, S., Crovetto, B., Bucek, J., Nahass, R.G., Milano, M., and Brunetti, L. (2022). Comparative Efficacy of Early COVID-19 Monoclonal Antibody Therapies: A Retrospective Analysis. Open Forum Infect Dis 9, ofac080. 10.1093/ofid/ofac080.

40. Pinto, D., Park, Y.-J., Beltramello, M., Walls, A.C., Tortorici, M.A., Bianchi, S., Jaconi, S., Culap, K., Zatta, F., De Marco, A., et al. (2020). Cross-neutralization of SARS-CoV-2 by a human monoclonal SARS-CoV antibody. Nature 583, 290–295. 10.1038/s41586-020-2349-y.

41. He, Q., Wu, L., Xu, Z., Wang, X., Xie, Y., Chai, Y., Zheng, A., Zhou, J., Qiao, S., Huang, M., et al. (2023). An updated atlas of antibody evasion by SARS-CoV-2 Omicron sub-variants including BQ.1.1 and XBB. Cell Rep Med 4, 100991. 10.1016/j.xcrm.2023.100991.

42. Qu, P., Evans, J.P., Faraone, J.N., Zheng, Y.M., Carlin, C., Anghelina, M., Stevens, P., Fernandez, S., Jones, D., Lozanski, G., et al. (2023). Enhanced neutralization resistance of SARS-CoV-2 Omicron subvariants BQ.1, BQ.1.1, BA.4.6, BF.7, and BA.2.75.2. Cell Host Microbe 31, 9–17.e13. 10.1016/j.chom.2022.11.012.

43. Smith, D.J., Lapedes, A.S., de Jong, J.C., Bestebroer, T.M., Rimmelzwaan, G.F., Osterhaus, A.D.M.E., and Fouchier, R.A.M. (2004). Mapping the Antigenic and Genetic Evolution of Influenza Virus. Science 305, 371–376. doi:10.1126/science.1097211.

44. Chalkias, S., McGhee, N., Whatley, J.L., Essink, B., Brosz, A., Tomassini, J.E., Girard, B., Wu, K., Edwards, D.K., Nasir, A., et al. (2023). Safety and Immunogenicity of XBB.1.5-Containing mRNA Vaccines. medRxiv, 2023.2008.2022.23293434. 10.1101/2023.08.22.23293434.

45. Koch, J., Uckeley, Z.M., Doldan, P., Stanifer, M., Boulant, S., and Lozach, P.Y. (2021). TMPRSS2 expression dictates the entry route used by SARS-CoV-2 to infect host cells. Embo j 40, e107821. 10.15252/embj.2021107821.

46. Hui, K.P.Y., Ho, J.C.W., Cheung, M.C., Ng, K.C., Ching, R.H.H., Lai, K.L., Kam, T.T., Gu, H., Sit, K.Y., Hsin, M.K.Y., et al. (2022). SARS-CoV-2 Omicron variant replication in human bronchus and lung ex vivo. Nature 603, 715–720. 10.1038/s41586-022-04479-6.

47. Essalmani, R., Jain, J., Susan-Resiga, D., Andréo, U., Evagelidis, A., Derbali, R.M., Huynh, D.N., Dallaire, F., Laporte, M., Delpal, A., et al. (2022). Distinctive Roles of Furin and TMPRSS2 in SARS- CoV-2 Infectivity. J Virol 96, e0012822. 10.1128/jvi.00128-22.

48. Meng, B., Abdullahi, A., Ferreira, I.A.T.M., Goonawardane, N., Saito, A., Kimura, I., Yamasoba, D., Gerber, P.P., Fatihi, S., Rathore, S., et al. (2022). Altered TMPRSS2 usage by SARS-CoV-2 Omicron impacts infectivity and fusogenicity. Nature 603, 706–714. 10.1038/s41586-022-04474-x.

