## Supplemental Table 1 and Supplemental figure 1 for "Immune Evasion, Infectivity, and Fusogenicity of SARS-CoV-2 Omicron BA.2.86 and FLip Variants": Qu_BA.2.86_SA.pdf

| | Bivalent HCWs (n=14) | 3-dose HCWs (n=15) | XBB.1.5-Wave Patients (n=11) |
|---|---|---|---|
| **Age in Years at Sample Collection** [Median (Range)] | 36 (25-48) | 33 (26-61) | 53 (36-75) |
| **Gender** [n (% of Total)] | | | |
| Male | 8 (57.1%) | 10 (66.7%) | 7 (63.6%) |
| Female | 6 (42.9%) | 5 (33.3%) | 4 (36.4%) |
| **Sample Collection Window** | Dec. 2022- early Jan. 2023 | Aug. 2021- Feb. 2022 | Feb. 2023- July. 2023 |
| **Vaccine status** [n (% of Total)] | | | |
| Unvaccinated | na | na | 3 (27.3%) |
| 2-dose Janseen | na | na | 1 (9.1%) |
| 2-dose Moderna | na | na | 1 (9.1%) |
| 3-dose Moderna | na | 3 (20%) | 1 (9.1%) |
| 3-dose Pfizer | na | 12 (80%) | 2 (18.2%) |
| 2-dose Pfizer+1-dose Pfizer bivalent | 1 (7.1%) | na | na |
| 3-dose Moderna+1-dose Moderna bivalent | 12 (85.8%) | na | 1 (9.1%) |
| 4-dose Moderna+1-dose Pfizer bivalent | 1 (7.1%) | na | 2 (18.2%) |
| **Sample Collection Timing** [Median (Range)] | | | |
| Days post 3$^{rd}$ dose for Recipients of three doses | na | 40 (14-86) | na |
| Days post the bivalent dose for Recipients | 66 (23-108) | na | na |
| **COVID-19 positive** [n (% of Total)] | 10 (71.4%) | 0 (0 %) | (100 %) |
| Days before sample collection [Median (Range)] | 254 (50-994) | na | na |
| **Infected Variants** | | | |
| XBB1.5 | na | na | 8 (73%) |
| Undermined | na | na | 3 (27%) |

**Table S1: Demographic and sample collection information of vaccinated HCWs and XBB.1.5-Wave COVID-19 positive patients, related to Figure 2 and Figure S1.** Summary of serum sample information for HCWs that received 1 dose of the Pfizer or Moderna bivalent mRNA vaccine and for HCW who received 3 doses of monovalent mRNA vaccines. 3 of the 10 HCWs who had received the bivalent booster were infected by SARS-CoV-2 before the Omicron wave, whereas the left 7 were positive during the Omicron wave (6 infected about 5 months before and 1 infected around 1 month after the bivalent vaccination). Summary information for the XBB.1.5-Wave infected patients in Columbus, Ohio is also provided. na means "not applicable".

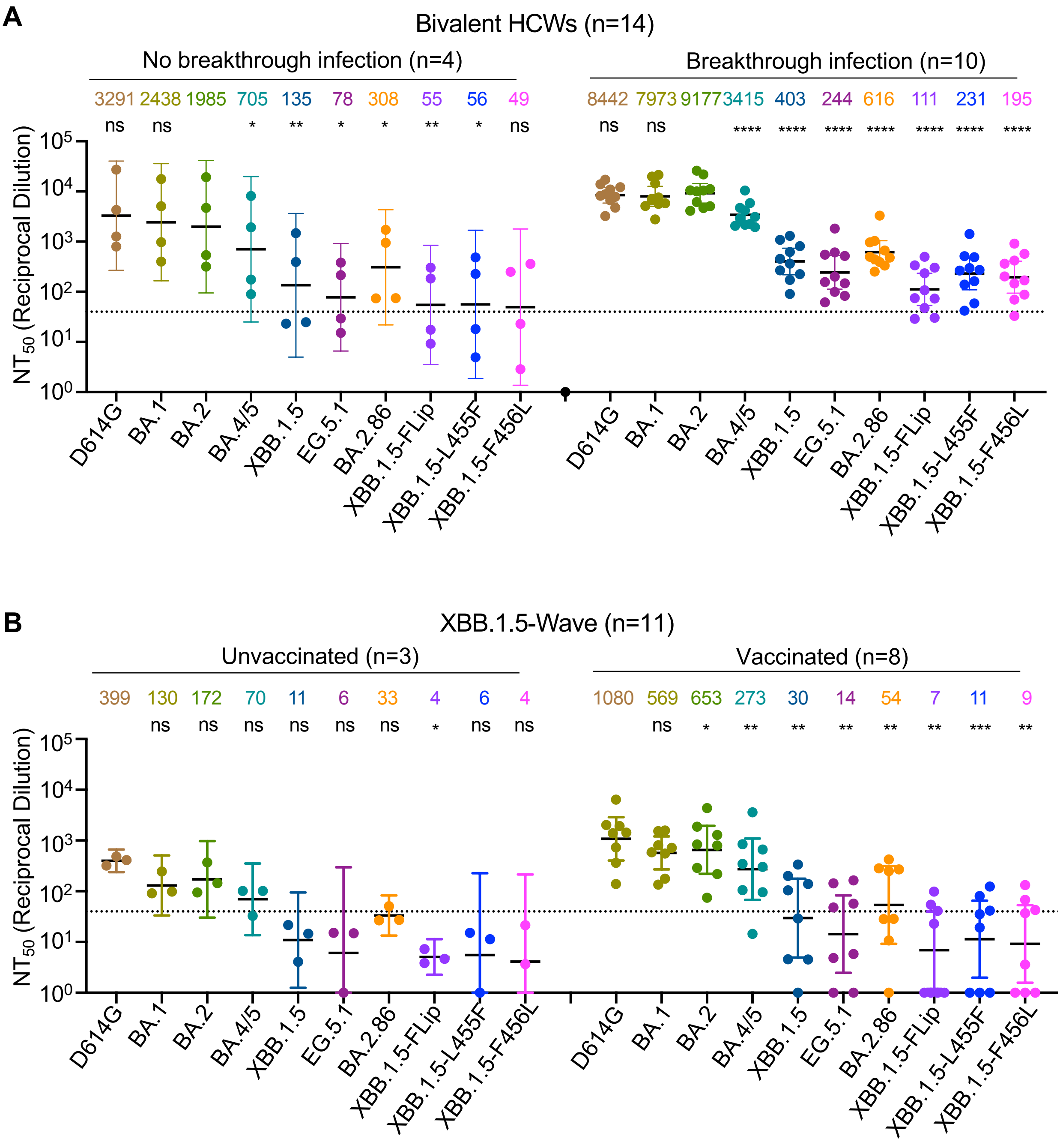


**Figure S1: Subgroup analyses of neutralization**. (**A**) $NT_{50}$ data shown in Figure 2A for bivalent HCWs sera were reanalyzed based on the HCWs' COVID-19 status, including no breakthrough infection (n = 4) and breakthrough infection (n = 10). (**B**) $NT_{50}$ data shown in Figure 2C for XBB.1.5-Wave sera is replotted by separating the vaccination status, i.e., Unvaccinated (n = 3) vs. Vaccinated (n = 8). Bars represent geometric means with 95% confidence intervals. Geometric mean $NT_{50}$ values are displayed on the top. Dashed lines represent the threshold of detection, i.e., NT50 of 40. P values are displayed as ns $p > 0.05$, *$p < 0.05$, **$p < 0.01$, ***$p < 0.001$, and ****$p < 0.0001$. Significance was determined by one-way repeated measures ANOVA using Bonferroni's multiple testing correction to make comparisons between multiple groups.

Figure S1
